# Acute or chronic depletion of macrophages in the dorsal root ganglia induces neuropathic pain after unilateral cervical spinal cord injury

**DOI:** 10.1101/2023.08.11.553038

**Authors:** Soha J. Chhaya, Jonathan Houston Richards, Grace A. Giddings, Megan Ryan Detloff

## Abstract

The inflammatory response at the spinal cord injury (SCI) epicenter and heightened macrophage presence in the dorsal root ganglia (DRG) has been well characterized after SCI and correlates with neuropathic pain. CCL2, a chemokine that acts as a macrophage chemoattractant and neuromodulator, is implicated in pain development, however, the role of the CCL2-CCR2 axis in the development of pain after SCI has not been explored. Here, we examined the role of CCL2-CCR2 signaling in macrophage recruitment to the DRG as well as the prolonged presence of macrophages in the DRG on the development and persistence of pain after SCI. Adult female Sprague-Dawley rats received a moderate, unilateral C5 contusion. Sandwich ELISA revealed that CCL2 is upregulated in the ipsilesional C7 and C8 DRGs in the first 24 hours post injury (hpi) and returns to naïve levels by 72 hpi. To prevent monocyte-derived macrophage recruitment to the DRG, additional SCI rats received vehicle or INCB3344, a CCR2 antagonist, intravenously at the time of SCI and at 24 and 48 hpi. INCB3344 administration induced transient forepaw allodynia at 7dpi in nearly all rats (88%) compared to only 33% in vehicle controls that resolves partially by 28 dpi, as measured by von Frey and mechanical conflict avoidance paradigms. As expected, qPCR analyses of whole DRG revealed that INCB3344 reduced macrophage markers and inflammatory cytokines in the ipsilesional C7 and C8 DRGs at 7 dpi compared to vehicle treated rats. By 28 dpi, there were no significant differences between INCB3344 or vehicle-treated groups, indicating that SCI-induced macrophage presence in the DRG is delayed by INCB3344 treatment. Moreover, gene expression of markers of macrophage polarity and cytokines suggest a pro-inflammatory environment in the DRG at 28dpi. DRG macrophage ablation via liposomal clodronate at 21dpi did not ameliorate hypersensitive pain behavior, though their ablation did reduce paw withdrawal thresholds in SCI rats that did not previously demonstrate pain behavior. Collectively, these data suggest that driving macrophages to a pro-reparative phenotype may be a viable and effective analgesic strategy that acts by modulating both the immune response and the experience of pain.

## Introduction

Monocyte-derived macrophages are involved in clearance of pathogens and debris through phagocytosis, and they also secrete cytokines and chemokines that mediate the function of adaptive and innate immune cells as well as neurons and ultimately behavior [32]. Chronic neuropathic pain correlates with a robust immune response in the dorsal horn of the spinal cord [81; 82; 95] [61; 103]. After spinal cord injury (SCI), similar inflammation occurs at the lesion site [26; 66] as well as at remote caudal and supraspinal [112] CNS regions involved in pain processing as well as in the dorsal root ganglia (DRG) of the peripheral nervous system [18].

After a neuropathic injury, such as SCI or peripheral nerve injury, damaged neurons and glial cells release the chemoattractant CCL2 as a result of a pro-inflammatory signaling cascade, leading to extravasation of monocytes from bone marrow and their recruitment to the damaged tissue to initiate inflammation [5; 11; 13; 21; 106; 110]. Additionally, CCL2 and other cytokines released in tandem in the DRG or dorsal horn of the spinal cord are pain mediators that can sensitize neurons and glial cells, leading to peripheral and central sensitization and chronic neuropathic pain [1; 9; 12; 20; 35; 67; 76; 90; 91; 97; 104; 107; 108; 115]. Macrophages recruited by CCL2 and other chemokines to sites of nerve injury or inflammation and along the sensory neuroaxis are involved in modulating the inflammatory environment. This neuroimmune response is now widely regarded as a critical component of neuropathic pain pathology [1; 3; 46; 48; 52; 54; 78; 80; 87; 93; 99; 102].

Recently, we have shown that a cervical spinal cord contusion induces an increase in the number of macrophages in the DRG that correspond to pain development in the forelimb innervated by primary sensory neurons at that level [18]. Previous work from our lab and others has elucidated the role of the immune response in SCI-induced pain including microglial/macrophage activation, astrogliosis and pro-inflammatory cytokine upregulation [25-27; 36; 38; 41; 42; 47; 63; 71; 75; 77; 86; 105; 113], however, these studies have mainly focused on the lesion epicenter or in the dorsal horns of the spinal cord. SCI-induced pain, however, has a complex etiology, involving dysfunction in primary sensory neurons in the DRG as well as pathological changes in supraspinal areas involved in pain [19; 39; 40; 62] where the role of neuroimmune interactions in generating sensory dysfunction has not been well studied. Additionally, the immune response after SCI is complex, and macrophages recruited to the lesion site as well as to other accessory sites can adopt pro-regenerative phenotypes critical to the wound healing response or inflammatory phenotypes that are neurotoxic [36]. Moreover, while CCL2 itself is secreted as a result of activation of pro-inflammatory pathways, the activation of CCL2-CCR2 signaling within the immune cells it recruits induces downstream anti-inflammatory pathways, leading to the release of anti-inflammatory cytokines and the modulation of macrophage and other immune cell to a more anti-inflammatory state [24; 83; 88].

Here, we tested the hypothesis that macrophages in the DRG are critical modulators of SCI-induced pain development and maintenance. To address the role of DRG macrophages in pain development, we pharmacologically inhibited early macrophage infiltration to the DRG by locally inhibiting the receptor for CCL2—CCR2— in the time window that CCL2 is upregulated. To address the role of DRG macrophages on established SCI-induced pain, we ablated DRG macrophages via intraganglionic liposomal clodronate three weeks after injury. We then assessed whether pain behavior or the immune environment in the DRG are modulated by macrophage manipulation. A portion of this data was reported in a doctoral dissertation [18].

## Materials and Methods

### Subjects

Adult, female Sprague-Dawley rats (225-250g, Charles River Laboratories) were housed two to three per cage on sawdust bedding in a controlled environment (12h light-12h dark cycle) with food and water *ad libitum*. All experimental procedures were approved by the Drexel University Institutional Animal Care and Use Committee. Rats were randomly assigned to naïve control (n=18) or SCI (n=121) groups. The SCI group was further subdivided into subgroups for CCL2 time course ELISA experiment (n=6/time point), pharmacological experiments to reduce early macrophage infiltration to the DRG (n=8 rats per group, 2 timepoints), and experiments to deplete macrophages once pain is established after SCI (n=9 per group). All rats were acclimated to handling for one or two 15-minute sessions/day for 7-10 days prior to surgery, behavior and/or sacrifice as applicable.

### Unilateral C5 spinal cord contusion surgeries

Rats in the SCI group were subjected to a moderate, unilateral spinal cord contusion injury at C5 as described previously (Detloff et al., 2012). This injury induces neuropathic pain development in approximately 40% of animals at chronic time points. Briefly, rats were anesthetized using a ketamine (60mg/kg)-xylazine (6mg/kg)-acepromazine (6mg/kg) (XAK) cocktail and a hemi-laminectomy was performed at C5, exposing the right dorsal surface of the spinal cord up to and partially over midline. The spinal column was stabilized in the Infinite Horizons device (Precision Systems and Instrumentation, Lexington, KY (Scheff et al., 2003)) and a custom probe was positioned 2 mm over the right dorsal surface of C5 spinal cord. The spinal cord and surgical field were submerged in sterile saline, and a contusion was performed with 200 kdyne force with 0 second dwell time, resulting in tissue displacement of 1500-1700 µm that was consistent across rats and groups (Table 1). The incision was closed in layers. Antibiotic (cefazolin, 160 mg/kg) and 5cc of lactated Ringer’s solution were administered subcutaneously at the time of surgery. Animals were monitored daily post-injury for signs of pain and distress until sacrifice.

### Intravenous injection of INCB3344

INCB3344 (MedChemExpress, Monmouth, NJ), a selective CCR2 antagonist (Brodmerkel et al., 2005), or vehicle (1.15% DMSO in sterile saline) was administered intravenously as described in (Van Steenwinckel et al., 2015). Briefly, INCB3344 was dissolved in vehicle to make a stock solution of 1mg/ml. To combat low solubility and prevent precipitation, the solution was kept in a water bath at 37°C for 30 minutes and/or sonicated as necessary. 100 μl of a 0.18mM solution (∼0.103mg/ml) of INCB3344 or vehicle was injected into the lateral tail vein of rats at the time of SCI, followed by additional i.v. injections at 24 hour and 48 hours post-SCI. The latter two injections were carried out under isoflurane anesthesia with an initial 5% isoflurane in O_2_ dose, followed by a 2% maintenance dose during injection to reduce restraint stress on the rat.

### Microinjection of liposomal clodronate into C7 and C8 DRGs

At 21 days post-SCI, 9 SCI rats demonstrating pain behaviors and 8 SCI rats with normal pain responses received intraganglionic injection of Clodrosome (Clod) into the C7 and C8 DRGs ipsilateral to the C5 SCI (Encapsula NanoSciences, Brentwood, TN). Briefly, rats were anesthetized with an XAK cocktail, and ipsilateral DRGs were exposed using a lateral incision and removal of the articular and transverse processes over C7 and C8. The tip of a glass micropipette attached to a Hamilton syringe was inserted 0.1mm into each DRG and 2ul of Clodrosome or Encapsome was injected at a rate of 0.2ul/min using a microinjection syringe pump (World Precision Instruments, Sarasota, FL). To prevent reflux of the Clodrosome or Encapsome solution, the micropipette was left in place for an additional 5 minutes prior to removal from the DRG. The incision was closed in layers. Cefazolin and lactated Ringer’s solution were administered post-operatively as described for spinal cord contusion surgeries, and rats were monitored daily for signs of pain and distress.

### Behavioral Testing

All behavioral testing was conducted as described in (Chhaya et al., 2019). Rats in the INCB, Clodronate, and associated vehicle-treated groups, as well as rats with 7dpi survival assigned to CCL2 ELISA were assessed for development of mechanical pain using von Frey test. All rats were acclimated to the apparatus prior to testing, and forelimb paw withdrawal thresholds recorded preoperatively as the baseline response, and weekly post-operatively until experimental endpoints in a blinded experimental design. Monofilaments of varying bending forces were applied to the ipsilesional forepaw beginning with the 15 g monofilament in accordance with the up-down method first described by Chaplan et al. [16]. The paw withdrawal threshold was determined as the force that elicited a paw withdrawal response in at least 50% of its applications and thresholds were represented as the percent of baseline paw withdrawal thresholds at each timepoint.

Operant testing for pain was performed in the fourth week post-SCI using the Mechanical Conflict-Avoidance Paradigm (MCAP; Noldus Technologies, Leesburg, VA,[18; 43]. Briefly, rats became adept at escaping an aversively bright chamber by crossing a runway to enter a dark chamber on 3 training sessions prior to test day. For the conflict-avoidance test, a series of nociceptive probes was introduced on to the runway floor at a height of 2 mm, and the latency to exit the aversive light chamber was measured.

### CCL2 ELISA

C7 and C8 DRGs of naïve animals and the C7 and C8 DRGs ipsilateral to the C5 SCI at 12, 24, 72,120 hours post injury (hpi) and 7 dpi were rapidly isolated from rats sacrificed with an overdose of Euthasol (390mg/kg of sodium pentobarbital and 50 mg/kg of phenytoin, intraperitoneally). Ipsilesional C7-8 DRGs of each animal were pooled together, were flash frozen in supercooled isopentane and stored at -80°C until use. At the time of analysis, tissue was weighed, suspended in RIPA buffer according to weight (100mg/ml) and homogenized on ice using a sonicator (Fisher Scientific, Sonic Dismembrator Model 100). Homogenate was centrifuged at 14000rpm for 40 minutes at 4°C and supernatant collected. Protein assay was performed using the BCA method (Pierce BCA Protein Assay kit, Thermo Fisher Scientific) to determine total protein concentration. CCL2/MCP-1 levels in DRG tissue lysate were assayed using an ELISA kit (Abcam, MCP-1(CCL2) Rat ELISA kit #ab100777) as per the protocol and reported as pg/mg total protein.

### qPCR

DRGs from naïve rats and SCI rats treated with i.v. INCB3344, intraganlionic clodronate, and respective vehicle controls were dissected as indicated for ELISAs. Pooled ipsilesional C7 and C8 DRGs of each animal were homogenized in 1ml RNA-Solv Reagent (EZNA Total RNA kit, Omega Bio-Tek, R-6834, Norcross, GA) with 20 ul 2-mercaptoethanol per 1 ml reagent. 100 ul chloroform was then added and sample centrifuged at 12,000 g at 4°C for 15 minutes to separate the aqueous and organic phase. The aqueous phase containing RNA was mixed with equal volume of 70% ethanol and RNA purification performed using HiBind RNA mini columns as per manufacturer’s instructions. The amount of total RNA was quantified using a NanoDrop 1000 spectrophotometer (Thermo Fisher Scientific, Wilmington, Delaware, USA). Purity was verified using the A60/A280 ratio for each sample as approximately 2 (between 1.9 and 2.1, ideally between 2.06 and 2.08). RNA was aliquoted and stored at -80°C until use. 1 ug of RNA was reverse transcribed into cDNA using the qScript XLT cDNA Supermix (QuantaBio, 95048, Beverly, MA) in the Eppendorf Mastercycler (Eppendorf, Hauppauge, NY) under the following conditions: 10 minutes at 25°C, 60 minutes at 42°C, 5 minutes at 85°C. cDNA was aliquoted and stored at -20°C.

The PCR reactions were performed according to MIQE guidelines using the StepOnePlus RealTime PCR system (Applied Biosystems, Thermo Fisher Scientific, Waltham, MA). Reactions were performed with a total volume of 25 µl: 12.5ul PerfeCTa SYBR Green Fast Mix (QuantaBio, 95-72 Beverly, MA), 1ul diluted cDNA (1:50), 1.5ul each of 2.5uM forward and reverse primers and 8.5ul nuclease-free water with the following conditions: 50 °C for 2 min, 95 °C for 10 min, followed by 40 cycles at 95 °C for 15 s, and 60 °C for 1 min. All samples were run in triplicates and outliers reported by the StepOne Plus software were removed from consideration. Melt curve analysis was performed (95 °C for 15 s, and 60 °C for 1 min, 95 °C for 15 s) and no-template controls were run to verify absence of primer-dimers. GAPDH and 18s were used as the reference genes, their values averaged, and results were analyzed using the comparative cycle threshold (ΔΔCt) method and divided by average naïve ΔΔCt value to derive fold change relative to naive. All primer sequence information is available in Table 2. Primers were designed using PrimerBLAST engine (http://www-ncbi-nlm-nih-gov.ezproxy2.library.drexel.edu/tools/primer-blast/index.cgi, specificity verified using BLASTn (National Center for Biotechnology Information, National Institutes of Health, Bethesda, MD, (http://blast.ncbi.nlm.nih.gov/Blast.cgi?PROGRAM=blastn&PAGE_TYPE=BlastSearch&LINK_LOC=blasthome) and synthesized by Integrated DNA Technologies (Coralville, Iowa). Primer sequences were tested over 5-6 log dilutions of cDNA and standard curve generated to verify efficiency of primer (90-105%, R^2^=0.99).

### Statistics

All statistical analyses were performed in Sigmplot 13. Data are presented as mean ± SEM. All data sets were tested for normality with the Shapiro–Wilk test. Normally distributed data were tested with parametric tests: the t test or 1-way analysis of variance (ANOVA) followed by the Tukey test for each pair-wise comparison. Data that were not normally distributed were tested with nonparametric tests: ANOVA on Ranks, the Mann–Whitney U followed by the Dunn test for each pair-wise comparison. Statistical significance was set at p≤0.05, and all reported values are 2-tailed.

## Results

### SCI parameters are consistent across groups

SCIs were verified as consistent across groups and timepoints. There were no significant differences in the following metrics: force of the probe hit, displacement of cord upon impact, and velocity of the probe (Table 1).

### Transient elevation of CCL2 in ipsilesional DRG at early timepoints post-SCI

CCL2 is a macrophage chemoattractant protein that is responsible for extravasation of monocytes and their recruitment to tissue. CCL2 protein levels were elevated at 12 (p=0.0092) and 24 hpi (p=0.0127) in the C7 and C8 DRGs ipsilesionally after SCI (Fig 1 A, n=6/group). CCL2 protein levels were not significantly different from naïve by 7dpi, which is the earliest that allodynia is detectable in the forelimb after SCI. Fifty percent (8/16) rats demonstrated allodynia at 7dpi, defined as a reduction of more than 30% from baseline paw withdrawal thresholds (Fig 1 B). CCL2 levels were not significantly different between rats with and without allodynia at 7dpi (Fig 1 C).

**Figure 1.**
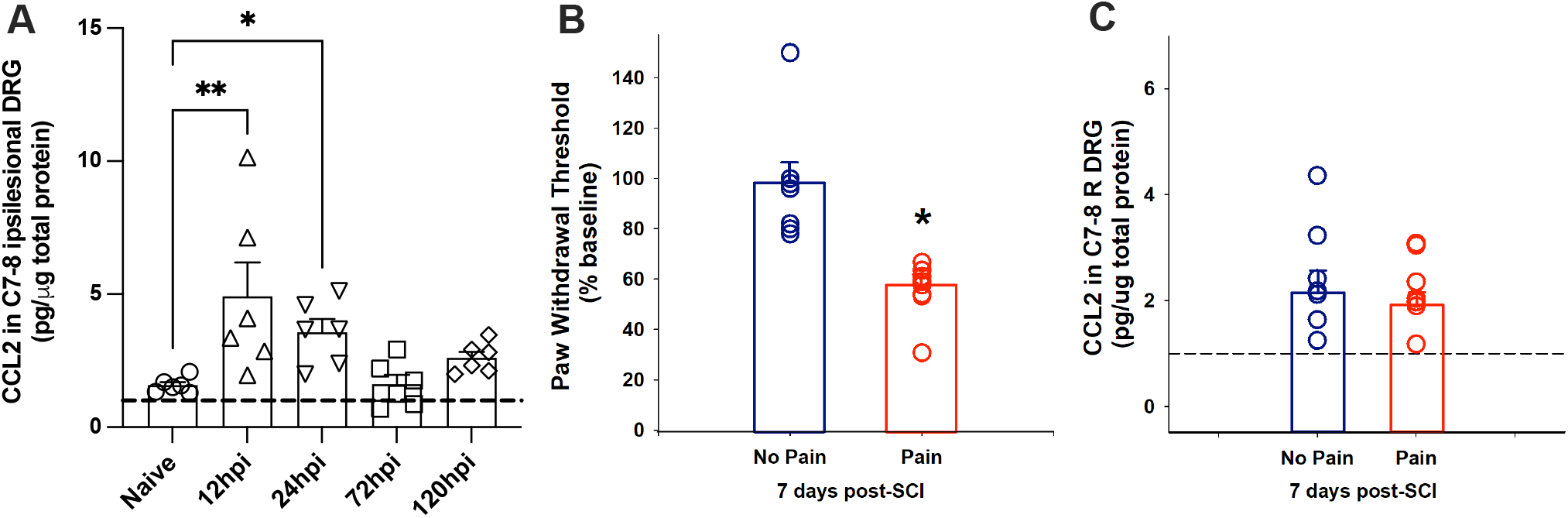
Increase in CCL2 protein in C7-8 ipsilesional dorsal root ganglia (DRG) early after spinal cord injury (SCI). CCL2 protein is significantly elevated in the ipsilesional DRG at 12- and 24-hours post-injury (hpi) compared to naïve rats (*p=0.0092, 12hpi vs naïve, *p=0.0127 24hpi vs naïve) and is near naïve-levels by 72 hpi to 120 hpi (p>0.05) (**A**, pg/ug total protein, n=6 per time-point). (8/16) rats demonstrated allodynia at 7 dpi, defined as a reduction of more than 30% from baseline paw withdrawal thresholds **(B)**. CCL2 remains near-naïve levels at 7days post-injury (dpi) and is not different in rats that do or do not have pain at that time point **(C)**. Dashed line indicates naïve. Group averages ± SEM are represented as bars and individual data points as open circles.

### Increased macrophage presence in the DRG at sub-acute and chronic timepoints post-SCI

We quantified changes in mRNA levels of CCL2 and its receptor CCR2 at 7dpi (vehicle n=17, INCB3344 n=16) and 28dpi (vehicle n=11, INCB3344 n=13) after treatment with a CCR2 antagonist, INCB3344 in 1.15% DMSO and saline, or vehicle, intravenously at the time of SCI and until 3dpi. CCL2 mRNA was not elevated at 7dpi in either vehicle- or INCB3344-treated rats, in fact, it was significantly downregulated at 7dpi (p<0.001, Fig 2 A), which is corroborated by lack of elevation in CCL2 protein at that time point (Fig 1). It remained downregulated at 28 dpi (p<0.001, Fig 2B). CCR2 mRNA was elevated at 7dpi in both treatment groups (p<0.05, Fig 2C), suggesting that CCR2+ cells are present in the DRG regardless of INCB3344 treatment. It also indicates that other cells that express CCR2 may be increasing their expression in response to SCI, at least at 7dpi when pain is first detectable. CCR2 was also elevated at 28dpi in vehicle- and INCB3344-treated rats (p<0.05, Fig 2D), indicating possible lasting presence of CCR2-recruited cells at chronic time points post-SCI, or a second phase of recruitment beyond the early infiltration of CCR2+ cells from the periphery.

**Figure 2.**
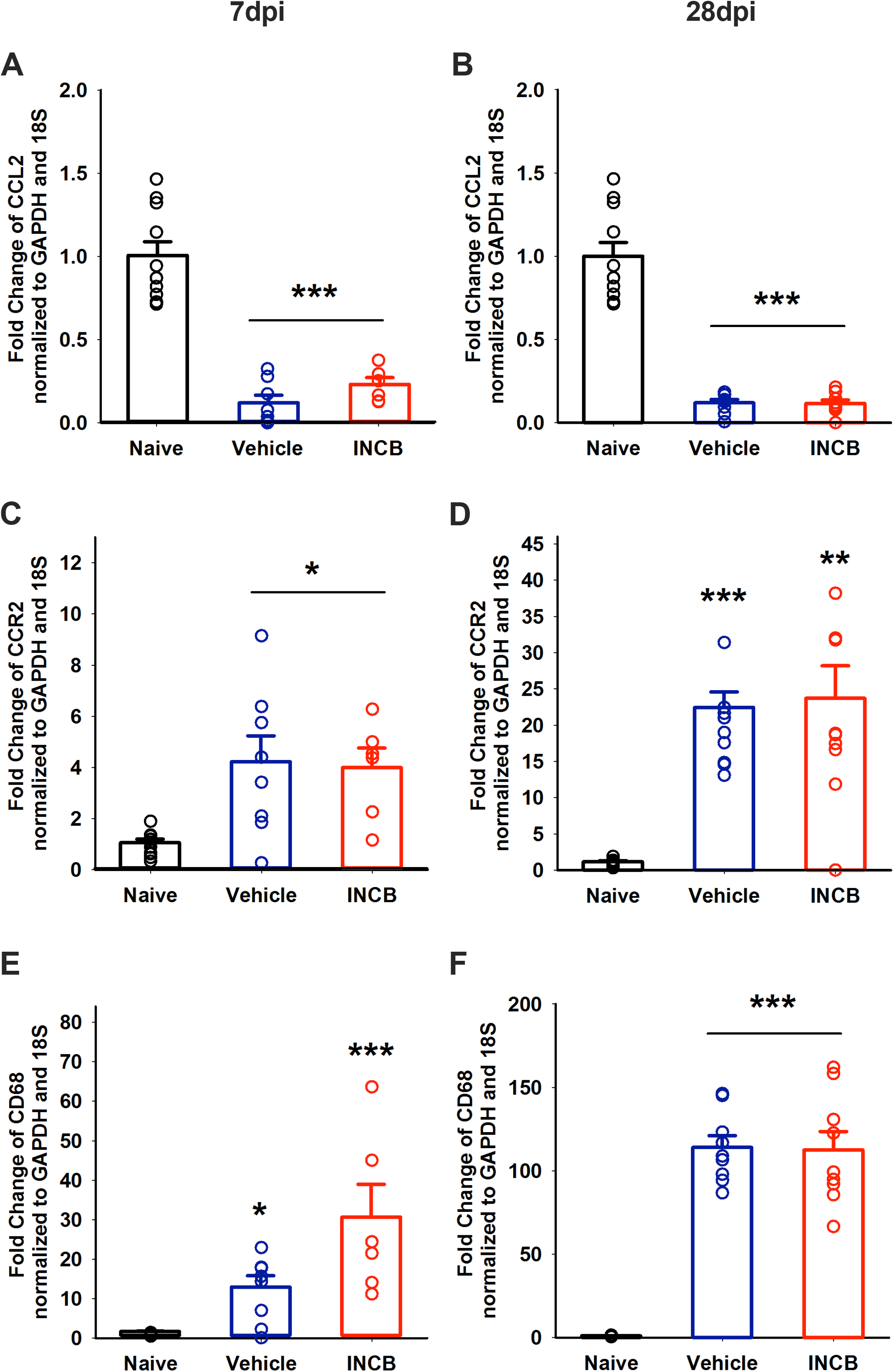
CCL2, CCR2 and CD68 expression at 7 and 28dpi in the C7-8 ipsilesional dorsal root ganglia (DRG) normalized to housekeeping genes GAPDH and 18S in vehicle (n=17 7dpi, n=11 28dpi)- and INCB3344-treated (n=16 7dpi, n=13) rats, expressed as fold change over expression in naïve rats. CCL2 mRNA is significantly downregulated by 7 dpi in vehicle- and INCB3344-treated rats (***p<0.001, compared to naive) **(A)** and remains downregulated at 28dpi (***p<0.001 compared to naïve) **(B)**. CCR2 mRNA is significantly elevated at 7dpi **(C)** in both vehicle- and INCB3344-treated rats (*p=0.021 vehicle compared to naïve, *p=0.026 INCB3344 compared to naïve) and at 28 dpi **(D)** in both treatment groups (***p=0.001 vehicle compared to naïve, **p=0.004 INCB3344 compared to naive). By 7 dpi the expression of CD68 mRNA in the DRG **(E)** is significantly increased in vehicle (*p=0.039)- and INCB-treated rats (***p<0.001) compared to naive. At 28 dpi CD68 mRNA expression **(F)** is similarly significantly elevated for vehicle- and INCB-treated rats (***p<0.001 compared to naïve). Group averages ± SEM are represented as bars and individual data points as open circles.

By 7dpi, CD68 mRNA was significantly elevated (p<0.05, Fig 2E), regardless of INCB3344 treatment. CD68 is expressed by phagocytic macrophages, and its elevation indicates presence of these cells by 7 dpi despite inhibition of recruitment for the first 3 dpi. At 28dpi, CD68 mRNA was significantly upregulated in both treatment groups (p<0.001, Fig 2F), consistent with previous findings of CD68+ cells in DRG tissue at chronic timepoint post-SCI[18].

### Development of transient allodynia when early macrophage infiltration is prevented by i.v. INCB3344 treatment

We administered INCB3344 intravenously in rats immediately after SCI until 3dpi, and assessed pain behavior preoperatively and weekly after SCI. At 7dpi, the vehicle-treated rats (n=8) had average paw withdrawal thresholds at 96.35 ±16.29% of baseline (p>0.05), while INCB3344-treated rats (n=9) had thresholds at 55.41±7.64% of baseline (p=0.0018, Fig 3 A). 33% of rats in the vehicle group demonstrated a reduction in paw withdrawal threshold of >50% from baseline, compared to 88% of rats in the INCB3344-treated group. By 28dpi, paw withdrawal thresholds for vehicle-treated rats and INCB3344-treated rats were not significantly different from naïve (p=0.02) (Fig 3 B). 27% of rats in the vehicle treated group demonstrated allodynia, compared to 46% in the INCB3344 treatment group. We also assessed cognitive perception of pain using the mechanical conflict paradigm at 28dpi in both treatment groups after 3 days of pre-training. There was a non-significant increase in latency to exit the aversively-lit chamber in INCB3344-treated rats compared to vehicle-treated rats (Fig 3C, ns).

**Figure 3.**
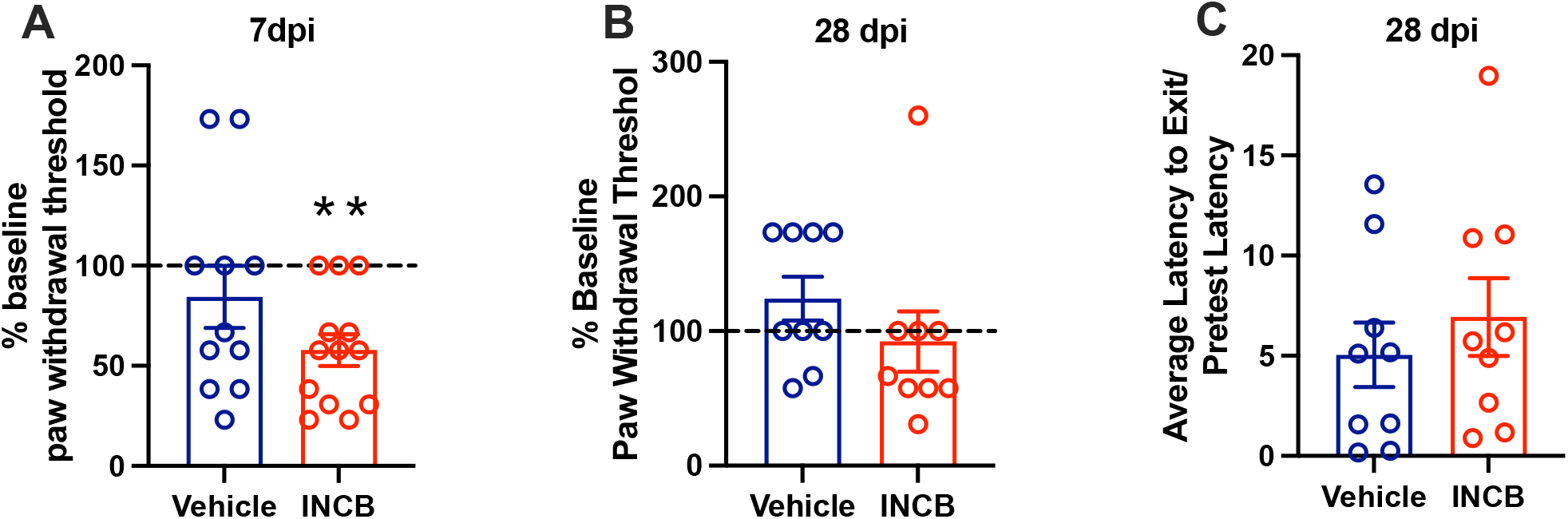
Assessment of evoked and ongoing pain behavior in rats treated with INCB3344 and vehicle-treated controls. Paw withdrawal thresholds were assessed preoperatively, 7 days and 28 days after spinal cord injury (SCI) in rats treated with vehicle (n=8) or INCB3344 (n=9) from the time of injury upto 3 days post-injury (dpi). INCB3344-treated rats had significantly lower paw withdrawal thresholds compared to vehicle-treated rats at 7 dpi (**p=0.0018, represented as percentage of baseline paw withdrawal threshold, **A**). By 28 dpi, there were no significant differences in paw withdrawal thresholds between vehicle- and INCB3344-treated rats (**B**). Cognitive perception of pain behavior was assessed at the experimental endpoint (28 dpi) using the mechanical conflict-avoidance paradigm. Rats were pre-trained in three training sessions to cross a runway to escape a brightly lit aversive chamber and get to a dark chamber. On testing day, the runway was inlaid with nociceptive probes and changes in latency to exit the bright chamber onto the nociceptive runway compared to pretest latency were measured. There were no significant differences in average latency to exit the aversively lit chamber compared to pretest latency to exit between the treatment groups. No rats in either group chose not to exit the aversively lit chamber (p=0.465). Group averages ± SEM are represented as bars and individual data points as open circles.

### mRNA expression of macrophage phenotype markers and cytokines in the DRG after SCI and vehicle or INCB3344 treatment

The inhibition of CCL2-CCR2 signaling acutely after SCI may influence the phenotype and secretome of macrophages recruited to the DRG after SCI, thereby modulating the microenvironment of sensory neurons in the DRG, affecting nociception. We analyzed mRNA expression of genes for two markers of a pro-inflammatory macrophage phenotype (iNOS and CD80), and two markers of anti-inflammatory macrophage phenotype markers (CD206 and Arg1) as well as several pro-inflammatory cytokines (tumor necrosis factor (TNF)-α, interleukin (IL)-6 and IL-12 and the anti-inflammatory cytokine IL-10 at 7dpi and 28dpi after intravenous administration of vehicle or INCB3344 from 0-3dpi.

Expression of the gene for pro-inflammatory macrophage marker CD80 was lower in INCB3344-treated rats compared to naïve and vehicle-treated rats at 7dpi (p<0.001, Fig 4A), but by 28dpi there is no significant difference in CD80 expression in treated rats compared to naïve rats (Fig 4B). However, there is evidence to suggest that pro-inflammatory macrophages exhibit a spectrum of markers and sub-states after SCI, and that CD80+ pro-inflammatory macrophages are not a dominantly expressed phenotype after SCI [36; 53]. Pro-inflammatory marker iNOS mRNA was significantly decreased at 7 dpi in INCB-treated rats compared to vehicle-treated or naïve rats (p=0.037 vehicle vs INCB, p=0.014 naïve vs INCB, Fig 4 C,), but by 28 dpi was elevated in both groups compared to naive (p<0.001, Fig 4 D).

**Figure 4.**
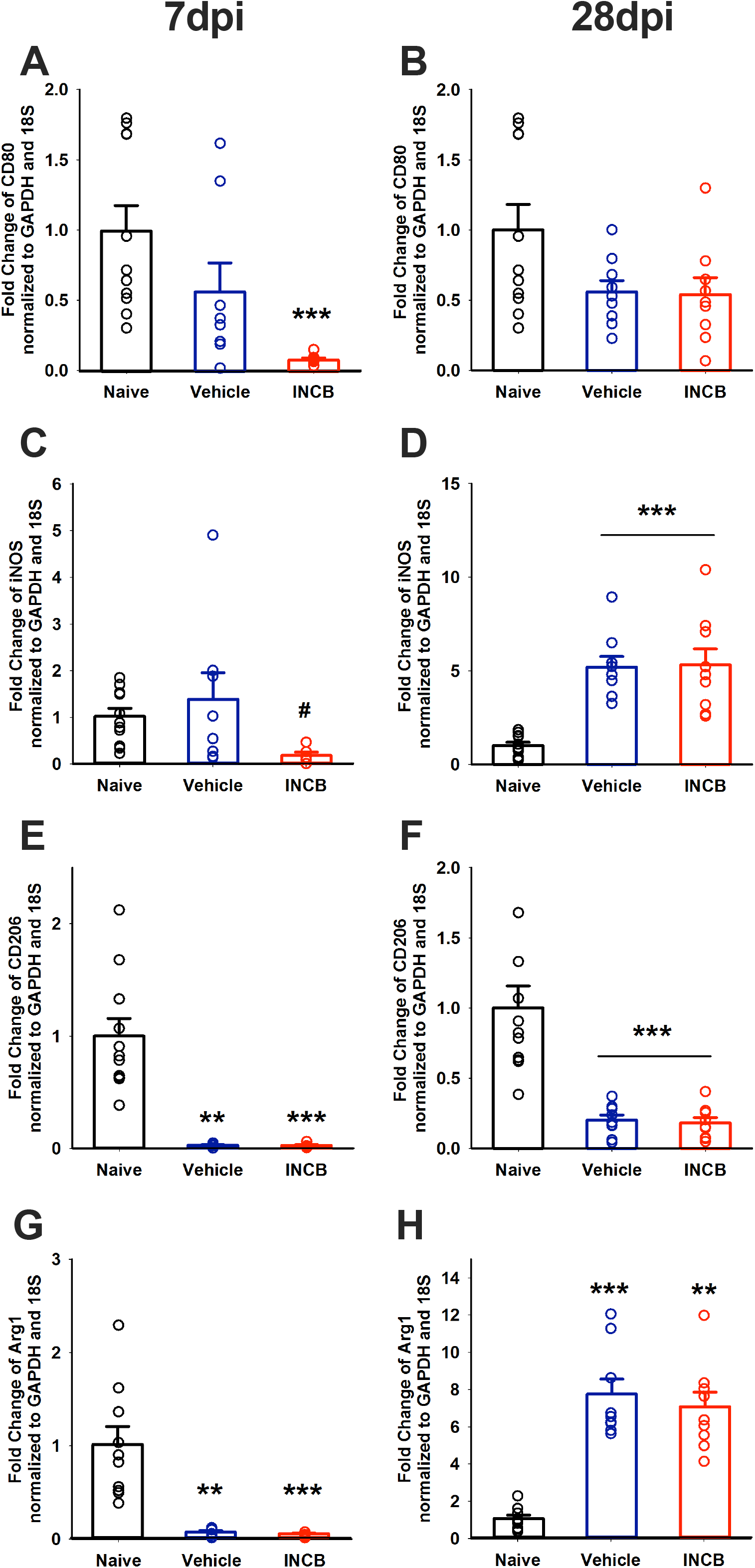
Expression of macrophage phenotype markers in the dorsal root ganglia after SCI with and without early macrophages. Expression of pro-inflammatory macrophage markers CD80 and iNOS, and anti-inflammatory CD206 and Arg1 mRNA normalized to housekeeping genes GAPDH and 18S in vehicle (n=17 7dpi, n=11 28dpi)- and INCB3344-treated (n=16 7dpi, n=13) rats, expressed as fold change over expression in naïve rats. At 7 dpi, CD80 is significantly decreased in INCB3344-treated rats compared to naive (***p<0.001, **A**). CD80 is not significantly elevated at 28 dpi in vehicle- or INCB3344-treated rats (**B**). iNOS is significantly decreased at 7 dpi in INCB3344-treated rats compared to naïve (#p=0.014) and vehicle-treated rats (#p=0.037) **(C)**. At 28 dpi, iNOS is significantly elevated in vehicle (***p<0.001) and INCB3344-treated rats compared to naïve (***p<0.001) (**D**). CD206 expression is significantly decreased after SCI at 7 dpi in vehicle (**p=0.002) and INCB3344-treated rats (***p<0.001) (**E**). By 28 dpi, CD206 is significantly decreased in both vehicle (***p<0.001) and INCB-treated rats (***p<0.001) compared to naïve (**F**). Arg1 is also significantly decreased at 7 dpi in both vehicle (**p=0.002)- and INCB3344-treated rats compared to naïve (***p=0.001) (**G**). Arg1 is elevated in vehicle-(***p<0.001) and INCB3344-treated rats (**p=0.002) at 28 dpi (**H**). Group averages ± SEM are represented as bars and individual data points as open circles.

Anti-inflammatory CD206 was expressed at significantly lower than naïve levels after SCI across treatment groups or time-points (p<0.001, Fig 4 E, F), and this may be because CD206 is only transiently elevated in anti-inflammatory M2a macrophages early after SCI and is downregulated in macrophages at the lesion at 28dpi [53]. At 7dpi, anti-inflammatory Arg1 expression was significantly decreased in both treatment groups compared to naïve (p=0.002 vehicle, p=0.001 INCB, Fig 4 G), but by 28dpi, while vehicle-treated rats had significantly elevated Arg1 expression, more so than INCB-treated rats (p<0.0001, Fig 4H), suggesting that transient INCB3344 treatment may be modulating DRG environment to then induce changes in inflammatory environment at later timepoints.

Gene expression levels of pro-inflammatory cytokine TNF at 7dpi and 28dpi were not significantly different from naïve for either treatment group (Fig 5 A, B). IL-6 expression was significantly increased at both 7 and 28 dpi, regardless of treatment (vehicle vs naïve p<0.001, INCB p=0.025 at 7dpi, p<0.001 vehicle vs naïve, p=0.012 INCB vs naïve, Fig 5C, D); IL-12 expression was significantly decreased after SCI at 7dpi in INCB3344-treated rats compared to naïve rats (p=0.025) and vehicle-treated rats (p=0.039) (Fig 5E), and significantly downregulated by 28dpi regardless of treatment (p<0.001, Fig 5F).

**Figure 5.**
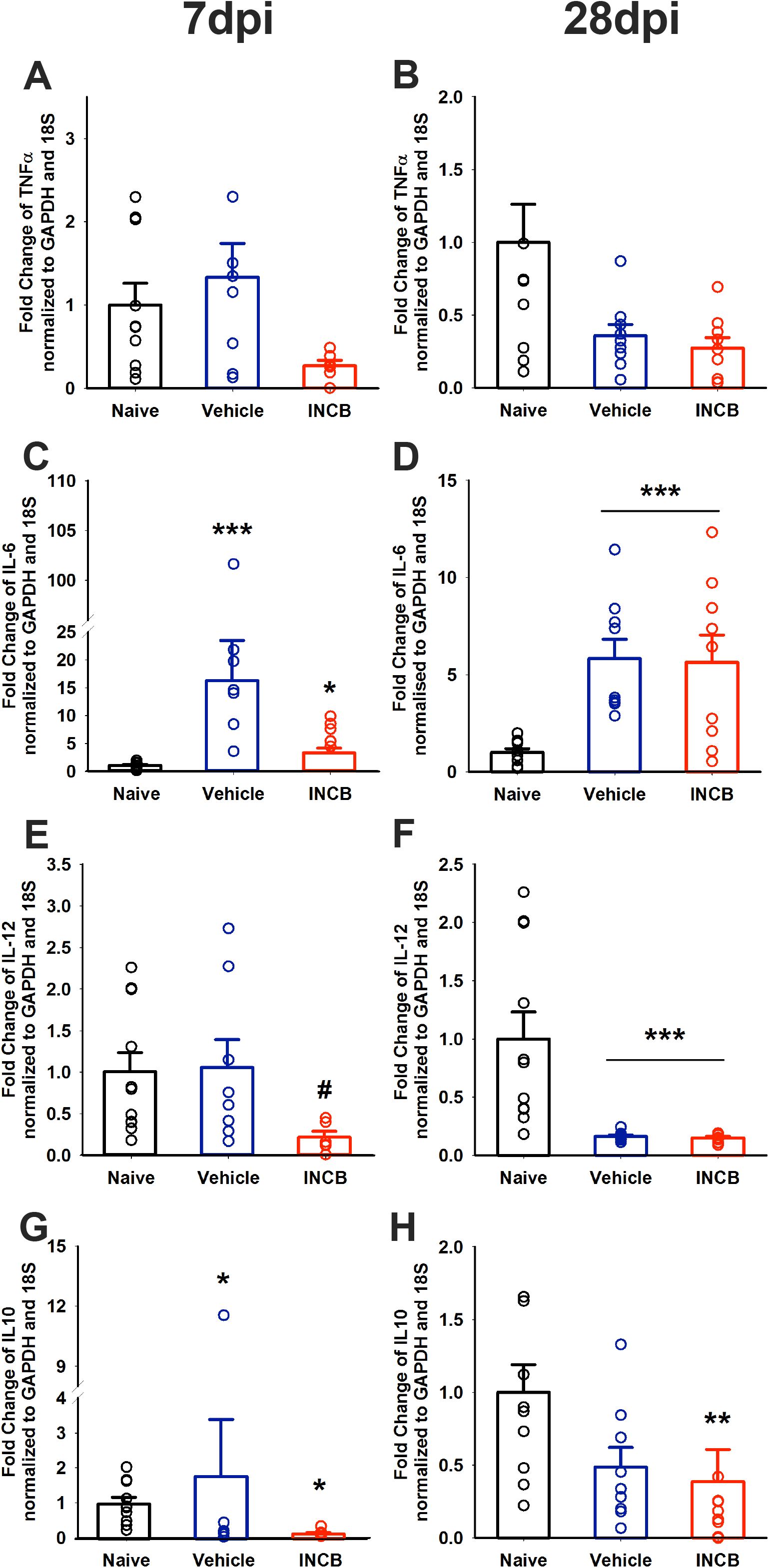
Expression of cytokines in the dorsal root ganglia following SCI and INCB3344 treatment. Expression of pro-inflammatory cytokines tumor necrosis factor α (TNFα), interleukin-6 (IL-6) and interleukin-12 (IL-12), and anti-inflammatory interleukin-10 (IL-10) mRNA normalized to housekeeping genes GAPDH and 18S in vehicle (n=17 7dpi, n=11 28dpi) and INCB3344-treated (n=16 7dpi, n=13) rats, expressed as fold change over expression in naïve rats. There is a non-significant decrease in TNFα at 7 dpi in INCB3344-treated rats compared to naïve rats (**A**). There is a non-significant decrease in TNFα at 28 dpi in both vehicle- and INCB3344-treated rats (**B**). IL6 is significantly elevated at 7 dpi in vehicle (***p<0.001) and INCB3344-treated rats (*p=0.025) compared to naïve (**C**) and is elevated at 28 dpi in both treatment groups compared to naïve (***p<0.001 vehicle vs naive, **p=0.012 INCB vs naive) (**D**). IL12 is significantly decreased in INCB3344-treated rats at 7 dpi compared to naïve (#p=0.025) and vehicle-treated rats (#p=0.039) (**E**) and is significantly decreased in both treatment groups at 28 dpi compared to naïve (***p<0.001) (**F**). IL10 is significantly increased in vehicle-treated rats at 7 dpi compared to naïve (*p=0.038) and significantly decreased in the INCB3344-treated rats compared to naïve (*p=0.011) (**G**) and is significantly decreased at 28dpi in INCB3344-treated rats compared to naïve (**p=0.010), but not vehicle-treated rats (**H**). Group averages ± SEM are represented as bars and individual data points as open circles.

Anti-inflammatory IL-10 is differentially expressed after SCI and INCB3344 treatment. At 7dpi, IL-10 was significantly increased in vehicle-treated (p=0.038) but significantly decreased in INCB-treated rats (p=0.011) compared to naïve rats (Fig 5G), and at 28dpi, IL-10 expression was significantly decreased in INCB3344-treated rats compared to naïve (p=0.010), but not in vehicle-treated rats (Fig 5H). This corroborates the decrease in Arg1 expression at 28dpi, and indicates that INCB3344 treatment leads to a delayed decrease in anti-inflammatory macrophages and cytokines after SCI.

### Development of allodynia when DRG macrophages are depleted one month after SCI

To test whether chronic SCI-induced pain is due to the presence of macrophages in the DRG at chronic time points, we administered liposomal clodronate or empty liposomes directly to the DRG after pain is established, at 21 days post-SCI and measured paw hypersensitivity 7 days later. Clodronate-encapsulating liposomes are engulphed by phagocytic cells, such as macrophages, and clodronate initiates their depletion via apoptotic pathways. Based on a broad database of efficacy of clodronate in depleting macrophages in different organs and routes of administration[34; 49; 70; 84], and we can assume that at 5-7 days post-clodronate, when behavior was tested, there are few, if any, macrophages in the DRG. Prior to injection, there were no significant differences in pain thresholds between groups of rats assigned to vehicle or clodronate treatment groups (data not shown, p=0.670), verifying that there was no bias in the distribution of rats to the two treatment groups. Roughly half of the vehicle treated group and half of the clodronate treated group demonstrated pain behavior prior to treatment.

Intraganglionic injection of empty liposome vehicle, did not significantly alter forepaw sensitivity in SCI rats that did not have pain prior to vehicle injection (p>0.05, Fig 6A, A’). Surprisingly, empty liposome administration in SCI rats with established pain exhibited significantly increased paw withdrawal thresholds, recovering to their pre-injury paw sensitivity (p=0.0056, Fig 6B, 6B’). These data suggest that even slight disruption of the DRG can alter paw sensitivity.

**Figure 6.**
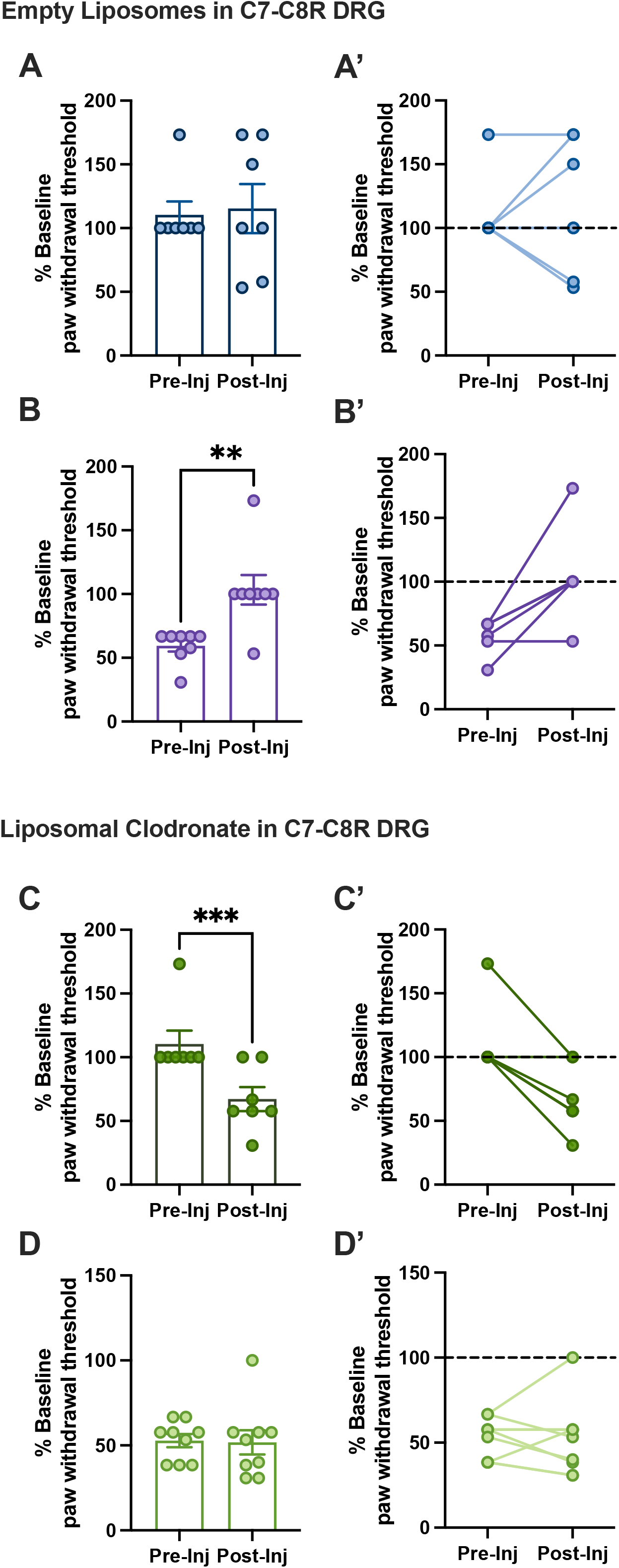
Effects of macrophage ablation on established SCI Pain. Injection of liposomal clodronate (n=17) or empty liposomes (n=14) in the DRG after SCI modulates withdrawal thresholds measured by the von Frey test using the up-down method. Injection of liposomal clodronate or its vehicle induced differing responses in rats depending on their pain state. There is no significant change in paw withdrawal thresholds (expressed as percent of baseline) before and 7 days after injection of empty liposomes in the C7 C8 right DRGs of rats that maintain normal pain thresholds after SCI (n=7, **A,** p>0.05, **A’** pre- and post-injection thresholds matched A’). There is a significant increase in paw withdrawal thresholds when empty liposomes are injected into DRGs of rats with SCI pain (n=8, p=0.0118, **B, B’**). When clodronate liposomes are injected into DRGs of rats without SCI pain, there is a significant decrease in paw withdrawal thresholds 7 dats post-injection (n=9, p=0.0001, **C, C’**). In rats with SCI pain prior to clodronate injection, there is no significant change in paw withdrawal thresholds before and 7 days after injection (n=7, p>0.05, **D, D’**). Group average± SEM are represented as bars and individual data points as filled circles.

On the other hand, macrophage ablation was unable to reverse existing pain, but induced pain in previously pain-free rats. Intraganglionic clodronate administration had no effect on paw withdrawal thresholds in SCI rats that exhibited allodynia before macrophage ablation (p=0.987, Fig 6D, D’). Interestingly, SCI rats that did not have pain prior to injection demonstrated a significant decrease in paw withdrawal thresholds with intraganglionic clodronate administration, (p=0.001, Fig 6C, C’). These data suggest that DRG macrophages in SCI rats with established pain and SCI rats with normal pain thresholds mat be different. Macrophages in SCI No Pain rats have a role in normalizing pain responses while macrophages in SCI Pain rats either are either ineffective at normalizing pain responses or are not involved in maintenance of established pain.

## DISCUSSION

Abundant literature suggests a detrimental role of macrophages and macrophage chemoattractant protein CCL2 in neuropathic pain development and persistence [1; 35; 91; 97; 106; 107; 111]. We have previously shown that macrophages are elevated in DRG at months-post SCI and correlate to the presence of pain [18]. Here, we provide evidence that macrophage function in the DRG, especially after SCI, is nuanced and complex with seemingly opposing roles at early and more chronic time points following spinal cord injury. We demonstrate that the monocyte chemoattractant CCL2 is significantly elevated in the DRG early and transiently following SCI. Surprisingly, acute inhibition of CCL2-CCR2 signaling after SCI induced neuropathic pain development, and despite a reduction in macrophage number, the cytokine milieu was skewed towards a more pro-inflammatory, deleterious state. In addition, depletion of DRG macrophages did not ameliorate established SCI pain, and macrophage depletion in pain-free SCI rats surprisingly induced paw hypersensitivity, despite the reduction in pan-macrophage markers with clodronate treatment. Collectively, these data underscore the premise that macrophage presence in the DRG may not be causative to sustained pain behavior.

While studies including Yu et al., [106] posit a deleterious role of DRG macrophages in pain initiation and persistence following a peripheral nerve injury, a growing body of work suggest a differential role of DRG macrophages [7; 98]. Further, several studies suggest that macrophage polarization states contribute to heterogeneous macrophage activities following peripheral or central injury [45; 65; 100]. For example, Van der Vlist et al., [98] demonstrate that M2a-like CD206+ macrophages ameliorate pain by donating mitochondria to nociceptors. As CD206+ macrophages are transiently active following injury, limiting macrophage infiltration at early time points impede their beneficial activity. Several studies have shown that macrophage polarization occurs in response to environmental signals [4; 53; 92; 101]. The environment of DRG neurons is particularly unique as they reside in both a central and peripheral environment, with cell bodies residing in the periphery and synapses in the central nervous system. This unique combination allows neurons to receive immune modulation at three distinct locations with differentially activated macrophages. Spinal cord injury produces only partial injury to primary afferents in the DRG, due in part to injury location, and the presence of Lissauer’s tract [44]. In this study, proprioceptive and tactile mechanoreceptor central axons which ascend ipsilaterally in the dorsal funiculus of the spinal cord are damaged, whereas primary nociceptors innervating the paw and located at spinal level C7-8, are spared. This combination of disrupted afferents produces a unique environment wherein spared nociceptors and injured proprioceptors intermingle [56]. Peripheral nerve injury, however, disrupts both afferent populations, which may result in a differential immune response. We posit that previous studies displaying differential responses to dampening CCL2 activity rely in part on the immune environment produced by peripheral rather than central injury.

Despite studies conducted in models of peripheral neuropathic pain demonstrating that depleting or knocking out macrophages in the DRG ameliorates pain behavior [17; 106], recent studies using macrophage-modulating therapies show no effect on pain behavior in models of SCI [55]. Moreover, Zhang et al., [109] demonstrated that delayed treatment with liposomal clodronate induced or worsened pain in a mouse model of lower back pain. While both peripheral and central mechanisms are involved in pain generation and maintenance, pain mechanisms may be different between peripherally- and centrally generated pain. The contribution of macrophages to pain pathology appears to be dependent on the type of pain and varies temporally, as demonstrated in this paper. Further, a key difference between the data presented here and previous research is that the inhibition is not through genetic knockout of a ligand, receptor, or cell type[2; 73]. Instead, the route of administration is peripheral (either intravenous or intraganglionic). These manipulations are not administered as a pre-treatment or continual treatment, but rather are delivered at key timepoints in the progression in the development and persistence of SCI-induced pain. These temporally controlled approaches allow for inferences to be drawn in an animal model that has targeted disruption of the naturally occurring immune response following SCI that more closely mimics DRG pathophysiology following human SCI.

Upon utilizing two interventions that influence the macrophage response at discrete time points post-SCI, we were able to investigate how the role of macrophages in the DRG changes as the injury evolves. Both acute and chronic macrophage inhibition led to an increase in pain behaviors in treated rats. Interestingly, the macrophage response to early inhibition via INCB3344 or liposomal clodronate ablation resulted in differential expression of macrophage markers 7 days later. Inhibition of early monocyte inhibition generated a delayed response or a presumptive rebound in macrophage presence as indicated by increased expression of macrophage markers. It can be inferred that this rebound in macrophage presence indicate that DRG macrophages play a protective role early after SCI.

Liposomal clodronate treatment led to depletion of DRG macrophages for the remainder of our experiment, and it can be inferred that DRG macrophages appear to be protective at chronic time points as well. Data from clodronate-treated SCI animals with normal nociception suggest that chronic macrophage presence in the DRG may be necessary for pain prevention. Importantly, rats with SCI pain injected with empty liposomes show pain resolution. The experimental design utilized in the clodronate experiments is reminiscent of the conditioning lesion paradigm in which a small peripheral nerve lesion preceding a central SCI leads to robust increases in axonal regeneration. While the order of the injuries is reversed in the current experiments, it is plausible that the succeeding intraganglionic injection of empty liposomes induces a typical early wound-healing and protective response whereby infiltrating macrophages may be predominantly anti-inflammatory and promote normative nociceptor functions survival [57; 74]. Regardless of the mechanism that underlies this reduction in paw hypersensitivity, the fact that such a small, localized perturbation in the DRG following SCI generated reductions in paw hypersensitivity is interesting and may have implications in chronic pain relief.

Importantly, SCI rats that maintained normal nociception did not develop pain upon injection of empty liposomes, further underlining our conclusions that macrophage influx alone may not be causative to pain. Their presence may indicate a yet to be fully understood communication network between neurons, resident glial cells, and immune cells in the DRG that drive plasticity of nociceptive and pain circuitry and ultimately changes in behavior [100]. Indeed, CCL2 is pleiotropic, and pharmacological inhibition of its receptor CCR2 is not specific to macrophage. CCR2 inhibition likely interrupts CCL2-CCR2 cascades between astrocytes, microglia, satellite glia and neurons [20; 76], and recently, satellite glia have been demonstrated to play a role in SCI Pain [22]. Also, resident glia in the DRG can be phagocytic under certain pathophysiological circumstances [68] and may also be targeted by liposomal clodronate. Therefore, it is possible that the effects of both INCB3344 and clodronate manipulations may be partially due to the disruption of the neuro-immune interactions between DRG nociceptors and glia or other immune cells [50]. To mitigate these limitations, our experiments were designed to specifically target macrophages prior to DRG invasion to minimize inhibitory effects of INCB3344 on resident DRG cells.

Damaged nociceptors release macrophage-modulating cytokines, and neuro-immune signaling cascades induce nociceptor dysfunction [14; 28; 29; 64; 72; 85]. The contribution of nociceptor hyperexcitability to central sensitization in SCI pain has recently been more thoroughly explored and certainly merits further investigation [6; 8; 15; 29]. Collectively, these data presented here support the longstanding theory that the development and maintenance of neuropathic pain have distinct neuroimmune mechanisms [23; 51; 102] and indicate that the evolution of the inflammatory response in the DRG is an important factor in the development and maintenance of pain [37; 59]. Dynamic interactions between macrophages and neurons are mediated by inflammatory cytokines and modulating the macrophage response temporally induced lasting effects on the inflammatory milieu in the DRG [96]. While inhibiting early monocyte recruitment with INCB3344 reduced the expression of both pro-inflammatory and anti-inflammatory macrophage markers and cytokines in the DRG early after SCI, it appears to have lasting effects on anti-inflammatory genes; IL-10 remains downregulated in the DRG at one month post SCI after INCB treatment. IL-10 has recently been studied in peripheral pain models, and its upregulation in the DRG is required for pain resolution[58]. Indeed, many have reported that IL-10 can downregulate several pro-inflammatory cytokines[10; 30; 33; 60; 69; 79; 89; 94; 114]. Modulating the inflammatory environment of the DRG in favor of pro-inflammation may be contributing to nociceptor dysfunction that is a driver of aberrant neuropathic pain. Propentofylline, a glial modulator, downregulates macrophage production of pro-inflammatory cytokines and merits further study as a therapy for SCI Pain [31]. Future experiments could examine this hypothesis by inducing anti-inflammatory macrophage polarization in the DRG, and may demonstrate that early modulators of macrophage polarity, such as exercise, are a reliable strategy for harnessing neuroimmune signaling in the DRG for pain prevention and modulation after SCI [18; 93].

In conclusion, the macrophage response in the dorsal root ganglia after spinal cord injury is heterogenous and vacillates temporally and can therefore have either beneficial regulatory or detrimental influences on nociceptor dysfunction and pain after SCI. Future studies exploring the differential role of macrophages in the DRG in pain initiation and persistence will help understand neuroimmune mechanisms of pain development. A careful understanding of the bidirectional communication between immune cells and nociceptors will be important for alleviating both developing and established SCI-induced pain.

## Acknowledgements

Financial support was contributed by the National Institutes of Health NINDS R01 and H.E.A.L. Initiative Supplement NS NIH NINDS R01 NS097880 (MRD). No potential competing interests was declared by the authors.

## Uncategorized References

[1] Abbadie C, Bhangoo S, Koninck YD, Malcangio M, Melik-Parsadaniantz S, Fletcher AW. Chemokines and pain mechanisms. Brain Research Reviews 2009.

[2] Abbadie C, Lindia JA, Cumiskey AM, Peterson LB, Mudgett JS, Bayne EK, DeMartino JA, MacIntyre DE, Forrest MJ. Impaired neuropathic pain responses in mice lacking the chemokine receptor CCR2. Proceedings of the National Academy of Sciences 2003.

[3] Austin PJ, Berglund AM, Siu S, Fiore NT, Gerke-Duncan MB, Ollerenshaw SL, Leigh SJ, Kunjan PA, Kang JWM, Keay KA. Evidence for a distinct neuro-immune signature in rats that develop behavioural disability after nerve injury. Journal of Neuroinflammation 2015.

[4] Avraham O, Feng R, Ewan EE, Rustenhoven J, Zhao G, Cavalli V. Profiling sensory neuron microenvironment after peripheral and central axon injury reveals key pathways for neural repair. Elife 2021;10.

[5] Bajetto A, Bonavia R, Barbero S, Florio T, Schettini G. Chemokines and their receptors in the central nervous system. Frontiers in neuroendocrinology 2001;22:147–184.

[6] Bavencoffe A, Li Y, Wu Z, Yang Q, Herrera J, Eileen JK, Walters ET, Dessauer CW. Persistent electrical activity in primary nociceptors after spinal cord injury is maintained by scaffolded adenylyl cyclase and protein kinase A and is associated with altered adenylyl cyclase regulation. Journal of Neuroscience 2016.

[7] Bavencoffe AG, Spence EA, Zhu MY, Garza-Carbajal A, Chu KE, Bloom OE, Dessauer CW, Walters ET. Macrophage Migration Inhibitory Factor (MIF) Makes Complex Contributions to Pain-Related Hyperactivity of Nociceptors after Spinal Cord Injury. J Neurosci 2022;42(27):5463–5480.

[8] Bedi SS, Yang Q, Crook RJ, Du J, Wu Z, Fishman HM, Grill RJ, Carlton SM, Walters ET. Chronic Spontaneous Activity Generated in the Somata of Primary Nociceptors Is Associated with Pain-Related Behavior after Spinal Cord Injury. Journal of Neuroscience 2010.

[9] Belkouch M, Dansereau M-A, Goazigo AR-L, Steenwinckel JV, Beaudet N, Chraib A, Melik-Parsadaniantz S, Philippe S. The Chemokine CCL2 Increases Nav1.8 Sodium Channel. The Journal of neuroscience 2011.

[10] Bethea JR, Nagashima H, Acosta MC, Briceno C, Gomez F, Marcillo AE, Loor K, Green J, Dietrich WD. Systemically Administered Interleukin-10 Reduces Tumor Necrosis Factor-Alpha Production and Significantly Improves Functional Recovery Following Traumatic Spinal Cord Injury in Rats. Journal of Neurotrauma 2009.

[11] Bianconi V, Sahebkar A, Stephen LA, Pirro M. The regulation and importance of monocyte chemoattractant protein-1. Current Opinion in Hematology 2018;25:44–51.

[12] Biber K, Erik B. Neuronal CC chemokines: the distinct roles of CCL21 and CCL2 in neuropathic pain. Frontiers in Cellular Neuroscience 2014.

[13] Bose S, Jungsook C. Role of chemokine CCL2 and its receptor CCR2 in neurodegenerative diseases. Archives of pharmacal research 2013;36:1039–1050.

[14] Carlton SM. Nociceptive primary afferents: They have a mind of their own. Journal of Physiology 2014.

[15] Carlton SM, Du J, Tan HY, Nesic O, Hargett GL, Bopp AC, Yamani A, Lin Q, Willis WD, Hulsebosch CE. Peripheral and central sensitization in remote spinal cord regions contribute to central neuropathic pain after spinal cord injury. Pain 2009.

[16] Chaplan SR, Bach FW, Pogrel JW, Chung JM, Yaksh TL. Quantitative assessment of tactile allodynia in the rat paw. Journal of neuroscience methods 1994;53:55–63.

[17] Chen O, Donnelly CR, Ji RR. Regulation of pain by neuro-immune interactions between macrophages and nociceptor sensory neurons. Curr Opin Neurobiol 2020;62:17–25.

[18] Chhaya SJ, Quiros-Molina D, Tamashiro-Orrego AD, Houlé JD, Detloff MR. Exercise-Induced Changes to the Macrophage Response in the Dorsal Root Ganglia Prevent Neuropathic Pain after Spinal Cord Injury. Journal of Neurotrauma 2019;36:877–890.

[19] Christensen MD, Hulsebosch CE. Chronic central pain after spinal cord injury. Journal of neurotrauma 1997;14:517–537.

[20] Clark AK, Old EA, Malcangio M. Neuropathic pain and cytokines: current perspectives. Journal of pain research 2013;6:803–814.

[21] Conductier G, Blondeau N, Guyon A, Nahon J-L, Carole R. The role of monocyte chemoattractant protein MCP1/CCL2 in neuroinflammatory diseases. Journal of neuroimmunology 2010;224:93–100.

[22] Corinne A. L-K, Ingves M, Henry KW, Shiao R, Collyer E, Tuszynski MH, Campana WM. Analysis of the behavioral, cellular and molecular characteristics of pain in severe rodent spinal cord injury. Experimental neurology 2016;278:91–104.

[23] Costigan M, Befort K, Karchewsk L, Robert SG, D’Urso D, Allchorne A, Sitarski J, Mannion JW, Pratt RE, Woolf CJ. Replicate high-density rat genome oligonucleotide microarrays reveal hundreds of regulated genes in the dorsal root ganglion after peripheral nerve injury. BMC Neuroscience 2002.

[24] Daly C, Barrett JR. Monocyte chemoattractant protein-1 (CCL2) in inflammatory disease and adaptive immunity: therapeutic opportunities and controversies. Microcirculation (New York, NY: 1994) 2003;10:247–257.

[25] David S, Antje K. Repertoire of microglial and macrophage responses after spinal cord injury. Nature reviews Neuroscience 2011;12:388–399.

[26] Detloff MR, Lesley CF, McGaughy V, Longbrake EE, Popovich PG, Basso DM. Remote activation of microglia and pro-inflammatory cytokines predict the onset and severity of below-level neuropathic pain after spinal cord injury in rats. Experimental Neurology 2008.

[27] Detloff MR, Quiros-Molina D, Amy SJ, Daggubati L, Nehlsen AD, Naqvi A, Ninan V, Vannix KN, McMullen M-K, Amin S, Ganzer PD, Houlé JD. Delayed Exercise Is Ineffective at Reversing Aberrant Nociceptive Afferent Plasticity or Neuropathic Pain After Spinal Cord Injury in Rats. Neurorehabilitation and neural repair 2016;30:685–700.

[28] Dubin AE, Patapoutian A. Nociceptors: The sensors of the pain pathway. Journal of Clinical Investigation 2010;120:3760–3772.

[29] Edgar TW. Nociceptors as chronic drivers of pain and hyperreflexia after spinal cord injury: An adaptive-maladaptive hyperfunctional state hypothesis. 2012;3:309.

[30] Eijkelkamp N, Steen-Louws C, Sarita AYH, Willemen HLDM, Prado J, Lafeber FPJG, Heijnen CJ, Hack CE, Roon JAGv, Kavelaars A. IL4-10 fusion protein is a Novel drug to Treat persistent inflammatory pain. Journal of Neuroscience 2016;36:7353–7363.

[31] Ellis A, Wieseler J, Favret J, Kirk WJ, Rice KC, Maier SF, Falci S, Watkins LR. Systemic administration of propentofylline, ibudilast, and (+)-naltrexone each reverses mechanical allodynia in a novel rat model of central neuropathic pain. The journal of pain: official journal of the American Pain Society 2014;15:407–421.

[32] Escudier B, Dorval T, Chaput N, André F, Caby MP, Novault S, Flament C, Leboulaire C, Borg C, Amigorena S, Boccaccio C, Bonnerot C, Dhellin O, Movassagh M, Piperno S, Robert C, Serra V, Valente N, Le Pecq JB, Spatz A, Lantz O, Tursz T, Angevin E, Zitvogel L. Vaccination of metastatic melanoma patients with autologous dendritic cell (DC) derived-exosomes: results of thefirst phase I clinical trial. J Transl Med 2005;3(1):10.

[33] Fenn AM, Hall JCE, Gensel JC, Popovich PG, Godbout JP. IL-4 signaling drives a unique arginase+/IL-1β+ microglia phenotype and recruits macrophages to the inflammatory CNS: Consequences of age-related deficits in IL-4Rα after traumatic spinal cord injury. Journal of Neuroscience 2014;34:8904–8917.

[34] Ferenbach DA, Sheldrake TA, Dhaliwal K, Kipari TMJ, Marson LP, Kluth DC, Hughes J. Macrophage/monocyte depletion by clodronate, but not diphtheria toxin, improves renal ischemia/reperfusion injury in mice. Kidney international 2012;82:928–933.

[35] Gao YJ, Ru Rong J. Chemokines, neuronal-glial interactions, and central processing of neuropathic pain. Pharmacology and Therapeutics 2010.

[36] Gensel JC, Zhang B. Macrophage activation and its role in repair and pathology after spinal cord injury. Brain research 2015;1619:1–11.

[37] Grace PM, Fabisiak TJ, Green-Fulgham SM, Anderson ND, Strand KA, Kwilasz AJ, Galer EL, Walker FR, Greenwood BN, Maier SF, Fleshner M, Watkins LR. Prior voluntary wheel running attenuates neuropathic pain. Pain 2016;157:2012–2023.

[38] Gwak YS, Kang J, Unabia GC, Hulsebosch CE. Spatial and temporal activation of spinal glial cells: Role of gliopathy in central neuropathic pain following spinal cord injury in rats. Experimental Neurology 2012;234:362–372.

[39] Hains BC, Saab CY, Waxman SG. Changes in electrophysiological properties and sodium channel Na v1.3 expression in thalamic neurons after spinal cord injury. Brain 2005.

[40] Hains BC, Saab CY, Waxman SG. Alterations in burst firing of thalamic VPL neurons and reversal by Na v1.3 antisense after spinal cord injury. Journal of Neurophysiology 2006.

[41] Hains BC, Waxman SG. Activated microglia contribute to the maintenance of chronic pain after spinal cord injury. Journal of Neuroscience 2006.

[42] Hansen CN, Norden DM, Faw TD, Deibert R, Wohleb ES, Sheridan JF, Godbout JP, Basso DM. Lumbar Myeloid Cell Trafficking into Locomotor Networks after Thoracic Spinal Cord Injury. Experimental neurology 2016;282:86–98.

[43] Harte SE, Meyers JB, Donahue RR, Taylor BK, Morrow TJ. Mechanical conflict system: A novel operant method for the assessment of nociceptive behavior. PLoS ONE 2016.

[44] Helgren ME, Goldberger ME. The recovery of postural reflexes and locomotion following low thoracic hemisection in adult cats involves compensation by undamaged primary afferent pathways. Exp Neurol 1993;123(1):17–34.

[45] Herd HL, Bartlett KT, Gustafson JA, McGill LD, Ghandehari H. Macrophage silica nanoparticle response is phenotypically dependent. Biomaterials 2015;53:574–582.

[46] Hu P, Alison LB, Keay KA, McLachlan EM. Immune cell involvement in dorsal root ganglia and spinal cord after chronic constriction or transection of the rat sciatic nerve. Brain, Behavior, and Immunity 2007.

[47] Hu X, Du L, Liu S, Lan Z, Zang K, Feng J, Zhao Y, Yang X, Xie Z, Wang PL, Ver Heul AM, Chen L, Samineni VK, Wang YQ, Lavine KJ, Gereau RWt, Wu GF, Hu H. A TRPV4-dependent neuroimmune axis in the spinal cord promotes neuropathic pain. J Clin Invest 2023;133(5).

[48] Iwai H, Ataka K, Suzuki H, Dhar A, Kuramoto E, Yamanaka A, Goto T. Tissue-resident M2 macrophages directly contact primary sensory neurons in the sensory ganglia after nerve injury. J Neuroinflammation 2021;18(1):227.

[49] Jackie EB, Enos RT, Velázquez KT, Carson MS, Nagarkatti M, Nagarkatti PS, Chatzistamou I, Davis JM, Carson JA, Robinson CM, Murphy EA. Macrophage depletion using clodronate liposomes decreases tumorigenesis and alters gut microbiota in the AOM/DSS mouse model of colon cancer. American journal of physiology Gastrointestinal and liver physiology 2018;314:G22–G31.

[50] Jain A, Hakim S, Woolf CJ. Unraveling the Plastic Peripheral Neuroimmune Interactome. J Immunol 2020;204(2):257–263.

[51] Ji RR, Chamessian A, Yu Qiu Z. Pain regulation by non-neuronal cells and inflammation. Science 2016.

[52] Jin X, Zheng W, Chi S, Cui T, He W. miR-506-3p Relieves Neuropathic Pain following Brachial Plexus Avulsion via Mitigating Microglial Activation through Targeting the CCL2-CCR2 Axis. Dev Neurosci 2023;45(1):37–52.

[53] Kigerl KA, Gensel JC, Ankeny DP, Alexander JK, Donnelly DJ, Popovich PG. Identification of two distinct macrophage subsets with divergent effects causing either neurotoxicity or regeneration in the injured mouse spinal cord. Journal of Neuroscience 2009.

[54] Kiguchi N, Kobayashi Y, Saika F, Sakaguchi H, Maeda T, Shiroh K. Peripheral interleukin-4 ameliorates inflammatory macrophage-dependent neuropathic pain. Pain 2015.

[55] Kopper TJ, McFarlane KE, Bailey WM, Orr MB, Zhang B, Gensel JC. Delayed Azithromycin Treatment Improves Recovery After Mouse Spinal Cord Injury. Front Cell Neurosci 2019;13:490.

[56] Kupari J, Usoskin D, Parisien M, Lou D, Hu Y, Fatt M, Lönnerberg P, Spångberg M, Eriksson B, Barkas N, Kharchenko PV, Loré K, Khoury S, Diatchenko L, Ernfors P. Single cell transcriptomics of primate sensory neurons identifies cell types associated with chronic pain. Nat Commun 2021;12(1):1510.

[57] Kwon MJ, Shin HY, Cui Y, Kim H, Thi AHL, Choi JY, Kim EY, Hwang DH, Byung Gon K. CCL2 mediates neuron-macrophage interactions to drive proregenerative macrophage activation following preconditioning injury. Journal of Neuroscience 2015.

[58] Laumet G, Bavencoffe A, Edralin JD, Huo XJ, Walters ET, Dantzer R, Heijnen CJ, Kavelaars A. Interleukin-10 resolves pain hypersensitivity induced by cisplatin by reversing sensory neuron hyperexcitability. Pain 2020;161(10):2344–2352.

[59] Laumet G, Edralin JD, Dantzer R, Heijnen CJ, Kavelaars A. Cisplatin educates CD8+ T cells to prevent and resolve chemotherapy-induced peripheral neuropathy in mice. Pain 2019;160(6):1459–1468.

[60] Ledeboer A, Brian MJ, Sloane EM, Mahoney JH, Langer SJ, Milligan ED, Martin D, Maier SF, Johnson KW, Leinwand LA, Chavez RA, Watkins LR. Intrathecal interleukin-10 gene therapy attenuates paclitaxel-induced mechanical allodynia and proinflammatory cytokine expression in dorsal root ganglia in rats. Brain, behavior, and immunity 2007;21:686–698.

[61] Lerch JK, Puga DA, Bloom O, Popovich PG. Glucocorticoids and macrophage migration inhibitory factor (MIF) are neuroendocrine modulators of inflammation and neuropathic pain after spinal cord injury. Semin Immunol 2014;26(5):409–414.

[62] Li C, Jonathan AS, Lowery-Gionta EG, McElligott ZA, McCall NM, Lopez AJ, McKlveen JM, Pleil KE, Kash TL. Mu Opioid Receptor Modulation of Dopamine Neurons in the Periaqueductal Gray/Dorsal Raphe: A Role in Regulation of Pain. Neuropsychopharmacology 2016;41:2122–2132.

[63] Li Z, Wei H, Piirainen S, Chen Z, Kalso E, Pertovaara A, Li T. Spinal versus brain microglial and macrophage activation traits determine the differential neuroinflammatory responses and analgesic effect of minocycline in chronic neuropathic pain. Brain, behavior, and immunity 2016;58:107–117.

[64] Lin Q, Wu J, William DW. Dorsal Root Reflexes and Cutaneous Neurogenic Inflammation After Intradermal Injection of Capsaicin in Rats. Journal of Neurophysiology 1999;82:2602–2611.

[65] Longbrake EE, Lai W, Ankeny DP, Popovich PG. Characterization and modeling of monocyte-derived macrophages after spinal cord injury. J Neurochem 2007;102(4):1083–1094.

[66] Lu X, Richardson PM. Responses of macrophages in rat dorsal root ganglia following peripheral nerve injury. J Neurocytol 1993;22(5):334–341.

[67] Luo P, Shao J, Jiao Y, Yu W, Weifang R. CC chemokine ligand 2 (CCL2) enhances TTX-sensitive sodium channel activity of primary afferent neurons in the complete Freud adjuvant-induced inflammatory pain model. Acta Biochimica et Biophysica Sinica 2018.

[68] Lutz AB, Chung W-S, Steven AS, Carson GA, Zhou L, Lovelett E, Posada S, Zuchero JB, Barres BA. Schwann cells use TAM receptor-mediated phagocytosis in addition to autophagy to clear myelin in a mouse model of nerve injury. Proceedings of the National Academy of Sciences of the United States of America 2017;114:E8072–E8080.

[69] Luzina IG, Keegan AD, Heller NM, Rook GAW, Shea-Donohue T, Atamas SP. Regulation of inflammation by interleukin-4: a review of “alternatives”. Journal of Leukocyte Biology 2012.

[70] Ma Y, Li Y, Jiang L, Wang L, Jiang Z, Wang Y, Zhang Z, Guo Yuan Y. Macrophage depletion reduced brain injury following middle cerebral artery occlusion in mice. Journal of Neuroinflammation 2016;13.

[71] Marchand F, Tsantoulas C, Singh D, Grist J, Anna KC, Bradbury EJ, McMahon SB. Effects of Etanercept and Minocycline in a rat model of spinal cord injury. European Journal of Pain 2009.

[72] Nascimento AI, Mar FM, Mónica Mendes S. The intriguing nature of dorsal root ganglion neurons: Linking structure with polarity and function. Progress in Neurobiology 2018.

[73] Nhong M, Wei T, Boring L, Israel FC, Ransohoff RM, Jakeman LB. Monocyte recruitment and myelin removal are delayed following spinal cord injury in mice with CCR2 chemokine receptor deletion. Journal of Neuroscience Research 2002;68:691–702.

[74] Niemi JP, DeFrancesco-Lisowitz A, Cregg JM, Howarth M, Zigmond RE. Overexpression of the monocyte chemokine CCL2 in dorsal root ganglion neurons causes a conditioning-like increase in neurite outgrowth and does so via a STAT3 dependent mechanism. Experimental Neurology 2016;275 Pt 1:25–37.

[75] Norden DM, Faw TD, McKim DB, Deibert RJ, Fisher LC, Sheridan JF, Godbout JP, Basso DM. Bone Marrow-Derived Monocytes Drive the Inflammatory Microenvironment in Local and Remote Regions after Thoracic Spinal Cord Injury. Journal of Neurotrauma 2019.

[76] Old EA, Marzia M. Chemokine mediated neuron-glia communication and aberrant signalling in neuropathic pain states. Current Opinion in Pharmacology 2012.

[77] Orr MB, Gensel JC. Spinal Cord Injury Scarring and Inflammation: Therapies Targeting Glial and Inflammatory Responses. Neurotherapeutics 2018.

[78] Parisien M, Lima LV, Dagostino C, El-Hachem N, Drury GL, Grant AV, Huising J, Verma V, Meloto CB, Silva JR, Dutra GGS, Markova T, Dang H, Tessier PA, Slade GD, Nackley AG, Ghasemlou N, Mogil JS, Allegri M, Diatchenko L. Acute inflammatory response via neutrophil activation protects against the development of chronic pain. Science Translational Medicine 2022;14(644):eabj9954.

[79] Park J, Joseph TD, Smith DR, Cummings BJ, Anderson AJ, Shea LD. Reducing inflammation through delivery of lentivirus encoding for anti-inflammatory cytokines attenuates neuropathic pain after spinal cord injury. Journal of Controlled Release 2018.

[80] Price TJ, Tadinada SM, Valtcheva MV, Krause EG, McIlvried LA, Dussor G, Kadunganattil S, Karlsson P, Usachev YM, Sheahan TD, Copits BA, Mickle AD, Haroutounian S, Mohapatra DP, Jain S, Shepherd AJ, Kloet ADd, Ray PR, Gereau RW. Angiotensin II Triggers Peripheral Macrophage-to-Sensory Neuron Redox Crosstalk to Elicit Pain. The Journal of Neuroscience 2018.

[81] Raghavendra V, Tanga F, DeLeo JA. Inhibition of microglial activation attenuates the development but not existing hypersensitivity in a rat model of neuropathy. J Pharmacol Exp Ther 2003;306(2):624–630.

[82] Raghavendra V, Tanga FY, DeLeo JA. Complete Freunds adjuvant-induced peripheral inflammation evokes glial activation and proinflammatory cytokine expression in the CNS. Eur J Neurosci 2004;20(2):467–473.

[83] Roca H, Zachary SV, Sud S, Craig MJ, Pienta KJ. CCL2 and interleukin-6 promote survival of human CD11b+ peripheral blood mononuclear cells and induce M2-type macrophage polarization. Journal of Biological Chemistry 2009.

[84] Rooijen Nv, Esther H. Liposomes for specific depletion of macrophages from organs and tissues. Methods in molecular biology (Clifton, NJ) 2010;605:189–203.

[85] Saleem M, Deal B, Nehl E, Jelena MJ, Pollock JA. Nanomedicine-driven neuropathic pain relief in a rat model is associated with macrophage polarity and mast cell activation. Acta Neuropathologica Communications 2019.

[86] Seiji O. The pathophysiological role of acute inflammation after spinal cord injury. Inflammation and Regeneration 2016.

[87] Shutov LP, Warwick CA, Shi X, Gnanasekaran A, Shepherd AJ, Mohapatra DP, Woodruff TM, Clark JD, Usachev YM. The Complement System Component C5a Produces Thermal Hyperalgesia via Macrophage-to-Nociceptor Signaling That Requires NGF and TRPV1. The Journal of neuroscience: the official journal of the Society for Neuroscience 2016;36:5055–5070.

[88] Sierra-Filardi E, Nieto C, Domínguez-Soto A, Barroso R, Sánchez-Mateos P, Puig-Kroger A, López-Bravo M, Joven J, Ardavín C, José LR-F, Sánchez-Torres C, Mellado M, Corbí AL. CCL2 shapes macrophage polarization by GM-CSF and M-CSF: idSierra-Filardi, E., Nieto, C., Domínguez-Soto, A., Barroso, R., Sánchez-Mateos, P., Puig-Kroger, A., Corbí, A. L. (2014). CCL2 shapes macrophage polarization by GM-CSF and M-CSF: identification. Journal of immunology (Baltimore, Md: 1950) 2014.

[89] Silva MDd, Bobinski F, Sato KL, Kolker SJ, Sluka KA, Santos ARS. IL-10 Cytokine Released from M2 Macrophages Is Crucial for Analgesic and Anti-inflammatory Effects of Acupuncture in a Model of Inflammatory Muscle Pain. Molecular Neurobiology 2014.

[90] Steenwinckel JV, Auvynet C, Sapienza A, Goazigo AR-L, Combadière C, Stéphane Melik P. Stromal cell-derived CCL2 drives neuropathic pain states through myeloid cell infiltration in injured nerve. Brain, Behavior, and Immunity 2015.

[91] Steenwinckel JV, Goazigo AR-L, Pommier B, Mauborgne A, Dansereau MAMA, Kitabgi P, Sarret P, Pohl M, Parsadaniantz SM, Goazigo ARL, Pommier B, Mauborgne A, Dansereau MAMA, Kitabgi P, Sarret P, Pohl M, Parsadaniantz SM. CCL2 released from neuronal synaptic vesicles in the spinal cord is a major mediator of local inflammation and pain after peripheral nerve injury. Journal of Neuroscience 2011.

[92] Stout RD, Suttles J. Functional plasticity of macrophages: reversible adaptation to changing microenvironments. Journal of Leukocyte Biology 2004;76(3):509–513.

[93] Sumizono M, Yoshizato Y, Yamamoto R, Imai T, Tani A, Nakanishi K, Nakakogawa T, Matsuoka T, Matsuzaki R, Tanaka T, Sakakima H. Mechanisms of Neuropathic Pain and Pain-Relieving Effects of Exercise Therapy in a Rat Neuropathic Pain Model. J Pain Res 2022;15:1925–1938.

[94] Szczepanik AM, Funes S, Petko W, Garth ER. IL-4, IL-10 and IL-13 modulate Aβ(1-42)-induced cytokine and chemokine production in primary murine microglia and a human monocyte cell line. Journal of Neuroimmunology 2001;113:49–62.

[95] Tanga FY, Raghavendra V, DeLeo JA. Quantitative real-time RT-PCR assessment of spinal microglial and astrocytic activation markers in a rat model of neuropathic pain. Neurochem Int 2004;45(2-3):397–407.

[96] Tansley S, Uttam S, Ureña Guzmán A, Yaqubi M, Pacis A, Parisien M, Deamond H, Wong C, Rabau O, Brown N, Haglund L, Ouellet J, Santaguida C, Ribeiro-da-Silva A, Tahmasebi S, Prager-Khoutorsky M, Ragoussis J, Zhang J, Salter MW, Diatchenko L, Healy LM, Mogil JS, Khoutorsky A. Single-cell RNA sequencing reveals time- and sex-specific responses of mouse spinal cord microglia to peripheral nerve injury and links ApoE to chronic pain. Nat Commun 2022;13(1):843.

[97] Thacker MA, Clark AK, Bishop T, Grist J, Yip PK, Moon LDF, Thompson SWN, Marchand F, McMahon SB. CCL2 is a key mediator of microglia activation in neuropathic pain states. European Journal of Pain 2009.

[98] van der Vlist M, Raoof R, Willemen H, Prado J, Versteeg S, Martin Gil C, Vos M, Lokhorst RE, Pasterkamp RJ, Kojima T, Karasuyama H, Khoury-Hanold W, Meyaard L, Eijkelkamp N. Macrophages transfer mitochondria to sensory neurons to resolve inflammatory pain. Neuron 2022;110(4):613–626.e619.

[99] Vega-Avelaira D, Sandrine MG, Fitzgerald M. Differential regulation of immune responses and macrophage/neuron interactions in the dorsal root ganglion in young and adult rats following nerve injury. Molecular pain 2009;5:70.

[100] Wang J, Wei Q, Yang Y, Che M, Ma Y, Peng L, Yu H, Shi H, He G, Wu R, Zeng T, Zeng X, Ma W. Small extracellular vesicles derived from four dimensional-culture of mesenchymal stem cells induce alternatively activated macrophages by upregulating IGFBP2/EGFR to attenuate inflammation in the spinal cord injury of rats. Front Bioeng Biotechnol 2023;11:1146981.

[101] Wang X, Cao K, Sun X, Chen Y, Duan Z, Sun L, Guo L, Bai P, Sun D, Fan J, He X, Young W, Ren Y. Macrophages in spinal cord injury: phenotypic and functional change from exposure to myelin debris. Glia 2015;63(4):635–651.

[102] Watkins LR, Maier SF. Beyond Neurons: Evidence That Immune and Glial Cells Contribute to Pathological Pain States. Physiological Reviews 2002.

[103] Winkelstein BA, DeLeo JA. Nerve root injury severity differentially modulates spinal glial activation in a rat lumbar radiculopathy model: considerations for persistent pain. Brain Res 2002;956(2):294–301.

[104] Wu XB, Zhu Q, Gao YJ. CCL2/CCR2 Contributes to the Altered Excitatory-inhibitory Synaptic Balance in the Nucleus Accumbens Shell Following Peripheral Nerve Injury-induced Neuropathic Pain. Neurosci Bull 2021;37(7):921–933.

[105] Xu J, Xiaoqiang E, Liu H, Li F, Cao Y, Tian J, Yan J. Tumor necrosis factor-alpha is a potential diagnostic biomarker for chronic neuropathic pain after spinal cord injury. Neuroscience Letters 2015.

[106] Yu X, Liu H, Hamel KA, Morvan MG, Yu S, Leff J, Guan Z, Braz JM, Basbaum AI. Dorsal root ganglion macrophages contribute to both the initiation and persistence of neuropathic pain. Nature Communications 2020;11(1):264.

[107] Zhang H, Jessica AB-D, Kosturakis AK, Li Y, Yoon S-Y, Walters ET, Dougherty PM. Induction of monocyte chemoattractant protein-1 (MCP-1) and its receptor CCR2 in primary sensory neurons contributes to paclitaxel-induced peripheral neuropathy. The journal of pain: official journal of the American Pain Society 2013;14:1031–1044.

[108] Zhang J, Shi XQ, Echeverry S, Mogil JS, Koninck YD, Rivest S. Expression of CCR2 in Both Resident and Bone Marrow-Derived Microglia Plays a Critical Role in Neuropathic Pain. Journal of Neuroscience 2007.

[109] Zhang L, Xie W, Zhang J, Shanahan H, Tonello R, Lee SH, Strong JA, Berta T, Zhang JM. Key role of CCR2-expressing macrophages in a mouse model of low back pain and radiculopathy. Brain Behav Immun 2021;91:556–567.

[110] Zhang X, Cheng J, Deng Y, Guo C, Cao Y, Wang S, Zhou C, Lin Z, Tang S, Zhou J. Identification and validation of biomarkers related to Th1 cell infiltration in neuropathic pain. J Inflamm (Lond) 2023;20(1):19.

[111] Zhang X, Wang YCaC, Huang LYM. Neuronal somatic ATP release triggers neuron-satellite glial cell communication in dorsal root ganglia. Proceedings of the National Academy of Sciences of the United States of America 2007.

[112] Zhao P, Waxman SG, Hains BC. Modulation of thalamic nociceptive processing after spinal cord injury through remote activation of thalamic microglia by cysteine-cysteine chemokine ligand 21. Journal of Neuroscience 2007.

[113] Zhou X, He X, Yi R. Function of microglia and macrophages in secondary damage after spinal cord injury. Neural regeneration research 2014;9:1787–1795.

[114] Zhou Z, Peng X, Hao S, Fink DJ, Mata M. HSV-mediated transfer of interleukin-10 reduces inflammatory pain through modulation of membrane tumor necrosis factor α in spinal cord microglia. Gene Therapy 2008.

[115] Zhu X, Cao S, Zhu MD, Liu JQ, Chen JJ, Yong Jing G. Contribution of chemokine CCL2/CCR2 signaling in the dorsal root ganglion and spinal cord to the maintenance of neuropathic pain in a rat model of lumbar disc herniation. Journal of Pain 2014.

